# Locus coeruleus injury modulates ventral midbrain neuroinflammation during DSS-induced colitis

**DOI:** 10.1101/2024.02.12.580010

**Authors:** Jake Sondag Boles, Jenny Holt, Cassandra L. Cole, Noelle K. Neighbarger, Nikhil M. Urs, Oihane Uriarte Huarte, Malú Gámez Tansey

## Abstract

Parkinson’s disease (PD) is characterized by a decades-long prodrome, consisting of a collection of non-motor symptoms that emerges prior to the motor manifestation of the disease. Of these non-motor symptoms, gastrointestinal dysfunction and deficits attributed to central norepinephrine (NE) loss, including mood changes and sleep disturbances, are frequent in the PD population and emerge early in the disease. Evidence is mounting that injury and inflammation in the gut and locus coeruleus (LC), respectively, underlie these symptoms, and the injury of these systems is central to the progression of PD. In this study, we generate a novel two-hit mouse model that captures both features, using dextran sulfate sodium (DSS) to induce gut inflammation and N-(2-chloroethyl)-N-ethyl-2-bromobenzylamine (DSP-4) to lesion the LC. We first confirmed the specificity of DSP-4 for central NE using neurochemical methods and fluorescence light-sheet microscopy of cleared tissue, and established that DSS-induced outcomes in the periphery, including weight loss, gross indices of gut injury and systemic inflammation, the loss of tight junction proteins in the colonic epithelium, and markers of colonic inflammation, were unaffected with DSP-4 pre-administration. We then measured alterations in neuroimmune gene expression in the ventral midbrain in response to DSS treatment alone as well as the extent to which prior LC injury modified this response. In this two-hit model we observed that DSS-induced colitis activates the expression of key cytokines and chemokines in the ventral midbrain only in the presence of LC injury and the typical DSS-associated neuroimmune is blunted by pre-LC lesioning with DSP-4. In all, this study supports the growing appreciation for the LC as neuroprotective against inflammation-induced brain injury and draws attention to the potential for NEergic interventions to exert disease-modifying effects under conditions where peripheral inflammation may compromise ventral midbrain dopaminergic neurons and increase the risk for development of PD.

## Introduction

Prodromal and early Parkinson’s disease (PD) is often marked by gastrointestinal distress and inflammation in the intestine. Constipation and defecatory dysfunction emerge in around 80%^1^ and 61%^2,3^, respectively, of people with PD, and it appears that gut dysfunction emerges years before a clinical diagnosis ^3–5^. Additionally, intestinal inflammation, reflected by greater fecal contents of pro-inflammatory cytokines^6^ and greater numbers of reactive glia in the colon^7^, and a deterioration of the colonic epithelium^8,9^ are evident in PD. And, people with inflammatory bowel disease (IBD) are at a roughly 22% greater risk of PD later in life^10^, and this risk is mitigated by the use of anti-TNF biologics^11,12^, further cementing a link between gut and brain in PD and highlighting inflammation as a potentially key mediator.

Studies in pre-clinical models have enabled investigators to directly investigate the extent to which gut inflammation can induce brain injury and inflammation. Multiple studies including several from our group have modeled IBD-like inflammation, using oral administration of dextran sulfate sodium (DSS) to denude the colonic epithelium^13^, and reported neuroimmune activation and immune cell infiltration in mice with colitis^14–18^. Critically, DSS-induced colitis has been reported to induce more robust nigrostriatal degeneration in mice expressing PD-relevant mutations in *LRRK2*^19,20^ and *SNCA*^21,22^ relative to their wild-type counterparts. While it is now widely accepted that PD arises from complex gene-by-environment interactions^23,24^, and mice with variants in PD-associated genes would be expected to display dysregulated responses to environmental insults, these mutations are overall low in frequency^24^. It was of greater interest to us to identify how gut inflammation might synergize with other frequent and early pathophysiological events in PD.

The disruption of norepinephrine (NE) signaling from the locus coeruleus (LC) is a common observation in PD and is believed to precede degeneration of the nigrostriatal pathway^25,26^. Emotional dysregulation, manifesting as clinical depression, anxiety, and/or apathy, and sleep disturbances, such as insomnia and daytime sleepiness, each emerge in 30-60% of people with PD^27–29^, and LC NE has been implicated in these non-motor PD symptoms^30,31^. A frank loss of LC NE neurons and decreased NE tone in the rest of the brain has been documented in people with PD with *in vivo* neuromelanin-sensitive MRI^32,33^, [^11^C]methyl-reboxetine PET imaging studies^34,35^ and *post mortem* immunohistochemical^36,37^ and biochemical^38^ studies. Interestingly, a similar pattern of global NE denervation has been observed in people with REM sleep behavior disorder (RBD)^39–41^, an established prodromal stage of PD or other synucleinopathies^42,43^, suggesting that LC injury occurs earlier than dopaminergic dysfunction. Indeed, mood dysfunction often emerges early and prior to a PD diagnosis^27,44,45^.

Critically, the death of the LC appears to be a central, disease-modifying process in PD. First, the non-motor symptoms described above seem to be associated with a worsened disease severity ^46,47^ and lower quality of life^48,49^. Second, studies in pre-clinical models have repeatedly demonstrated the neuroprotective function of the LC. A pharmacological lesion of the LC using N-(2-chloroethyl)-N-ethyl-2-bromobenzylamine (DSP-4) resulted in blood-brain barrier disruption^50^, neuroinflammation via the NF-κB inflammasome^51^, and oxidative stress^52^ in regions downstream from the LC. Pharmacological blockade of the β2-adrenergic receptor, simulating the loss of NE that occurs with LC degeneration, recapitulates the activation of neuroinflammation seen with LC neuron loss^53,54^. Treating primary microglial cells or organotypic brain slice cultures with NE or adrenergic receptor agonists further demonstrates the protective role of NE, with adrenergic receptor stimulation resulting in a suppression of microglial NF-κB, cytokine production, and motility^55,56^.

More interestingly, LC injury may interact with secondary disease-relevant insults to exacerbate neuroinflammation and neurodegeneration. A DSP-4 lesion has been shown to worsen proteinopathy and behavioral defects in mouse models over-expressing mutant tau^57^ and alpha-synuclein^58^. NE depletion through DSP-4 has also exacerbated nigrostriatal death and motor deficits of neurotoxin models of PD that selectively injury the midbrain dopaminergic neurons ^59–61^. Of most relevance to the gut-brain axis in PD, DSP-4 has exacerbated neuroinflammation after systemic inflammatory insults, including an acute lipopolysaccharide challenge^62–65^ and a high-sugar diet^66^. Clearly, the LC is an important regulator of neuroinflammation and brain health, especially amidst other inflammatory challenges, which would be expected to have important disease-modifying consequences on the development and progression of PD.

However, despite the frequency with which both emerge in PD, the role of LC death in neuroinflammation due to intestinal permeability has not yet been evaluated. Our hypothesis is that the LC protects the midbrain against leaky gut-induced neuroinflammation. Here, we pair the well-established DSS experimental colitis model with DSP-4, an LC-specific neurotoxin, to model the gut inflammation and LC injury that emerge frequently in PD and examine the combined effects of these insults on midbrain neuroimmune gene expression. Notably, we take care to affirm the specificity of this two-hit procedure, ensuring the NE deficit is restricted to the brain and the induction of colitis with DSS is unaffected. Specifically, based on relevant data using the DSP-4 model and other NE-related interventions and on our own data with the DSS-induced colitis model^14^, we expect that DSP-4-induced LC degeneration will exacerbate midbrain neuroimmune gene expression due to DSS-induced colitis, implicating an influential position for the LC as a protective locus against inflammatory stress along the gut-brain axis in PD. A better understanding of the relationship between gut health and LC permeability with respect to brain inflammation would advance our understanding of PD pathogenesis, emphasizing upstream disease-related events as potential diagnostic or therapeutic opportunities.

## Methods

### Animals

C57BL/6J mice were purchased from Jackson Laboratories (strain #000664), bred to congenicity, and housed until 2-3 months of age on a 12:12 light-dark cycle with *ad libitum* access to food and water. For experiments examining the specificity of DSP-4, mice were anesthetized with isoflurane and euthanized with transcardiac perfusion with heparinized PBS after their respective dosing schedules. Brains were rapidly extracted, and the frontal cortex was dissected using a razor blade, removing any olfactory bulb, striatum, or other tissue so cortical monoamines could be measured with high-performance liquid chromatography (HPLC). Colons were also extracted so roughly 5cm of tissue on the cecal end of the colon could be cleaned and analyzed with HPLC. For experiments employing a two-hit schedule with both DSP-4 and DSS, mice were again anesthetized and perfused as above, and colon lengths and spleen weights were measured. To assess colonic inflammation and injury, roughly 5cm of tissue on the cecal end of the colon was cleaned and flash-frozen in liquid nitrogen for biochemical and gene expression analyses. To assess neuroinflammation in this two-hit model, brains were extracted and the ventral midbrain was rapidly removed as described in ref. ^14^ and flash-frozen for gene expression analyses. All procedures were approved by the University of Florida Institutional Animal Care and Use Committee and followed the Guide for the Care and Use of Laboratory Animals from the National Institutes of Health.

### Drug administration

Within cages, mice were randomly assigned to DSP-4 or vehicle treatment groups. Mice were given a single intraperitoneal injection of DSP-4 (50mgkg) or sterile saline vehicle. DSP-4 (Sigma, #C8417) was dissolved in sterile saline to yield a 5mg/mL solution which was injected into mice. Due to the rapid half-life of DSP-4 in solutions with a physiological pH^67^, only enough solution was prepared for 3-5 mice at a time, and mice were injected within 2 minutes of the dissolution of DSP-4.

When combining DSP-4 with experimental colitis, cages were randomly assigned to receive dextran sulfate sodium (DSS) or untreated autoclaved tap water as a vehicle control. 40kDa DSS (ThermoFisher, #J14489-22) was dissolved in autoclaved tap water at the concentrations described below and placed in sterile water bottles for administration. To elicit peak intestinal injury, 3% DSS (w/v) was used for 7 days, and tissues were harvested. To elicit peak neuroinflammation, 2.5% DSS (w/v) was used for 7 days followed by 2 days of withdrawal with untreated autoclaved tap water. Our group has identified these schedules based on a previous study characterizing the kinetics of intestinal and brain inflammation in the DSS-induced colitis model^14^. During and after DSS administration, animal weights and disease severity were measured daily using the rubric described in Supplemental Table 1. In all two-hit studies, mice began receiving DSS treatment 28 days after a single injection of DSP-4 or vehicle.

### HPLC

Tissue samples were weighed and lysed in 0.1M perchloric acid. Colon samples were lysed in a 10:1 (volume to weight) ratio of 0.1M perchloric acid, while frontal cortex samples were lysed in a 20:1 ratio. Tissue was sonicated in pulses using a probe sonicator until tissue was fully dissociated. Tissue lysates were centrifuged at 16,000*g* for 15min at 4°C, the supernatants were filtered through a Nanosep 0.2µm mesh (Fisher Scientific, #50-197-9573) with centrifugation at 5,000*g* for 5min at 4°C, and samples were stored at –20°C until analysis. Monoamine concentrations were analyzed using high-performance liquid chromatography (HPLC) with electrochemical detection (HTEC, Azuma Inc.). A standard curve consisting of 3-methoxy-4-hydroxyphenylglycol (MHPG), NE, 3,4-dihydroxyphenylacetic acid (DOPAC), DA, 5-hydroxyindoleacetic acid (5HIAA), homovanillic acid (HVA), 3-methoxytyramine (3MT), and 5-hydroxytryptophan (5HT) were used to enable the measurement of these molecules at the picomolar level. Concentration data are shown as percentages of the control group.

### Intact tissue fluorescent labeling with iDISCO

Tissues were immunolabeled without sectioning using the iDISCO+ protocol without modifications^68,69^. Briefly, tissue was dehydrated with graded methanol baths, delipidated using a 2:1 solution of dichloromethane (DCM) and methanol, bleached in a solution of 3% hydrogen peroxide in methanol, and rehydrated through graded methanol baths. Tissue was permeabilized in a PBS buffer containing 2.3% (w/v) glycine, 20% dimethyl sulfoxide (DMSO), and 0.2% Triton X-100 for one day (colons, hindbrains) or two days (hemibrains) at 37°C. Non-specific binding was blocked in a PBS buffer containing 6% (w/v) normal donkey serum (NDS), 10% DMSO, and 0.2% Triton X-100 for one day (colons, hindbrains) or two days (hemibrains) at 37°C. Primary antibodies were applied in a PBS buffer containing 1000U/L heparin (Sigma, #H3393-50KU) and 0.2% Tween-20 (PTwH) plus 5% DMSO and 3% NDS at 37°C. Primary antibodies included rabbit IgG anti-norepinephrine transporter (Abcam, #ab254361), rabbit IgG anti-tyrosine hydroxylase (Millipore, #AB152), and rabbit IgG anti-dopamine beta-hydroxylase (Abcam, #209487). After 1-2 days of washes in PTwH, Alexa Flour 647-conjugated donkey anti-rabbit IgG antibody (Invitrogen, #A31573) was applied in PTwH and 3% NDS at 37°C. Antibody incubation times were 3 days for colon samples, 5 days for hindbrain samples, and 8 days for hemibrains, and all antibodies were used at a 1:500 dilution. Tissues were washed again for 1-2d in PTwH. At this point, colon samples were embedded in 1.2% agarose gel to facilitate mounting on a light-sheet microscope. Tissues were dehydrated in graded methanol baths, delipidated in a 2:1 solution of DCM and methanol, washed twice in pure DCM, and optically cleared by direct transfer to dibenzyl ether (DBE). Tissue remained in DBE without agitation until imaging.

### Fluorescence light-sheet microscopy

Cleared and immunolabeled intact tissues were imaged using an UltraMicroscope Blaze (Miltenyi Biotec). All images were acquired using right-side illumination, adaptive horizontal focus, 15% overlap between mosaic tiles, and a step-size half the width of the light-sheet for optimal axial resolution. Images were rendered in three dimensions using Imaris v. 9.8 (Oxford Instruments). Illumination settings, including laser power and exposure time, and image rendering settings, including any lookup table adjustments, were kept constant between images within an experiment.

### Nucleic acid and protein extraction

Protein and RNA were extracted from brain and colon using a TRIzol-based method. TRIzol (ThermoFisher, #15596018) was added to tissue and lysed using a sterile metal bead with rapid agitation in a TissueLyser II (Qiagen). Lysate was then mixed with UltraPure phenol:chloroform:isoamyl alcohol (ThermoFisher, #15593049) in a 5:1 ratio. Samples were centrifuged to separate protein, RNA, and DNA. The top aqueous layer containing RNA was transferred to a QIAshredder column (Qiagen, #79656), and flow-through as mixed 1:1 with 70% ethanol. This solution was transferred to a RNeasy Mini column (Qiagen, #74104) and centrifuged. The column was washed once in 700µL RW1 buffer and twice in 500µL RPE buffer, following the manufacturer’s instructions. RNA was eluted in nuclease-free water and a Denovix DS-11 spectrophotometer was used to measure nucleic acid concentrations.

To isolate protein, the white DNA layer and some of the pink layer was discarded, leaving roughly 300µL. 1mL of methanol was added to precipitate the protein. Protein precipitation occurred over 10 minutes at room temperature, and the protein was pelleted and washed in 500µL of methanol. The supernatant was decanted and the pellet was air-dried to remove residual methanol after which the protein was resuspended in 1% SDS (w/v) in deionized water. Protein concentrations were measured using a Pierce BCA Protein Assay Kit (ThermoFisher, #23225) according to manufacturer’s instructions.

### SDS-PAGE and immunoblotting

Within experiments, protein was diluted to an equivalent concentration for each sample in additional 1% SDS with 4X Laemmli sample buffer (Bio-Rad, #1610747) and 2-mercaptoethanol. 15µg of protein was loaded into a pre-cast 4-20% polyacrylamide Criterion TGX gel (Bio-Rad, #5678095). To normalize signal between different gels within an experiment, a constant sample was included on every gel. Electrophoresis consisted of one hour of 60V followed by 125V until finished, about 75 minutes. Electrophoresed samples were transferred to a PVDF membrane (Bio-Rad, #1620175) using a Trans-Blot Turbo Transfer system (Bio-Rad, #1704150) using the preset Mixed Molecular Weight program. Signal normalization was achieved using the Revert 700 Total Protein Stain (Li-Cor, #926-11011). The stain was imaged on a Li-Cor Odyssey Fc imager with a 2-minute exposure time.

After total protein visualization, the membrane was cut into sections containing proteins of interest and blocked in 5% (w/v) dry nonfat milk in TBS with 0.1% Tween-20 (TBST) for 1 hour at room temperature. Primary antibodies were applied overnight at 4°C in 5% milk (w/v) in TBST. Primary antibodies included rabbit IgG anti-ZO-1 (Invitrogen, #61-7300; 1:1000), rabbit IgG anti-occludin (Proteintech, #13409-1-AP; 1:5000), rabbit IgG anti-claudin-5 (Invitrogen, #34-1600, 1:1000), and rabbit IgG anti-NLRP3 (Cell Signaling Technologies, #15101S; 1:1000). After three washes in TBST, secondary goat IgG anti-rabbit IgG antibody conjugated to horseradish peroxidase (Jackson Immunoresearch, #111-035-144; 1:2000) was applied in 5% milk (w/v) in TBST for 1 hour at room temperature. Membranes were then washed three times with TBST and twice with TBS to remove detergent. SuperSignal West Pico (ThermoFisher, #A34580) or Femto (#A34095) were used, depending on the abundance of the targets, according to manufacturer’s instructions. These chemiluminescent reactions were imaged using a Li-Cor Odyssey Fc imager. Blot images were analyzed using ImageStudio Lite (Li-Cor), and target signal intensities were normalized to their respective Total Protein Image, which was then multiplied by a normalization factor derived from the ratio of normalized signal intensities between the constant samples on different blots.

### cDNA synthesis and qPCR

Complementary DNA (cDNA) was synthesized from RNA using the ImProm-II Reverse Transcription System (Promega, #A3800) according to manufacturer’s instructions. RNA was hybridized to oligo-dT primers in a total volume of 4µL for 5 minutes at 70°C. Then, the reverse transcription master mix, containing 1X ImProm-II reaction buffer, 5mM MgCl_2_, 0.6mM deoxyribonucleotides, RNase inhibitor, and reverse transcriptase, was added to the RNA with annealed primers. Samples were equilibrated at 25°C for 5 minutes followed by reverse transcription at 42°C for one hour and reverse transcriptase inactivation at 70°C for 15 minutes. Samples were diluted with nuclease-free water to 5-10µg/µL, depending on the abundance of the target gene.

Gene expression was quantified using quantitative polymerase chain reaction (qPCR) with SYBR Green chemistry. 5µL of diluted cDNA was mixed with 35µL of qPCR master mix, consisting of 1.25µM forward and reverse primers, SYBR Green (ThermoFisher, #A46112), and nuclease-free water. This reaction mix was plated into 10µL triplicates. A QuantStudio 5 Real-Time PCR System (ThermoFisher) was used to perform the qPCR. Triplicates were pruned based on intra-sample standard deviation and deviation from the median value using a custom R script. Data were analyzed using the ΔΔC_T_ method, with data being expressed as a fold change from the average ΔΔC_T_ of the reference group. Primer sequences can be found in Supplemental Table 2.

### NanoString nCounter gene expression analysis

To assess how our two-hit model impacts neuroimmune activation in the midbrain, we leveraged the nCounter direct transcript counting platform from NanoString Technologies using the Mouse Neuroinflammation Panel with an additional 55 genes added in that are known to be responsive to NE, affected by colitis, or involved in the immunological crosstalk between brain and periphery. Prior to running the assay, RNA quality was assessed using a Bioanalyzer 2100 (Agilent) with the RNA 6000 Nano Kit (Agilent, #5067-1511) according to manufacturer’s instructions. We used the 6 samples from each treatment group from both sexes with the highest RINs (48 samples total). From these samples, 100ng of total RNA was input into the nCounter protocol using the MAX/FLEX workstation, following manufacturer’s instructions.

Data quality was examined in R (v. 4.2-4.3) using the NACHO package^70^, which revealed two samples to be discarded ahead of downstream analyses (Fig. S1A-B). Inspecting the expression of individual housekeeping genes revealed the gene *Asb10* to be only rarely captured (Fig. S1C), so this gene was removed. Data were then normalized using the NanoTube package^71^ with the RUVg method from the RUVSeq package^72^, which uses only high-confidence housekeeper probes and removes unwanted technical variance while preserving biological variance. This strategy has been shown to outperform NanoString’s commercial normalization strategy, which uses only centrality measures from the housekeeper probes included in the panel^73^.

After normalization, differential expression testing was then performed with DESeq2^74^. Differentially expressed genes (DEGs) were classified as genes that showed | log_2_(fold change) | > 0.25 and Benjamini-Hochberg-adjusted *p*-value < 0.05 for a given comparison. DEGs relative to the H2O + saline control group were identified for each of the other treatment groups, collapsed across sex. Then, the lists of DEGs from the DSS + saline group were treated as co-expression modules for which the eigengenes were calculated for each sample using the WGCNA package^75^.

### Statistical analysis

All statistical analysis was performed in R v. 4.2-4.3. Tissue monoamine concentration changes due to DSP-4 were analyzed with a Wilcoxon rank sum test as data were not assumed to come from a normal distribution. Daily weight measurements during DSS were analyzed with an ANOVA that included day as a within-subjects variable and DSP-4 as a between-subjects variable. Sphericity of these data were assessed using the Performance package’s ‘check_sphericity’, which uses Mauchly’s test, and the Geisser-Greenhouse correction was applied when appropriate. Colon length, spleen weight, tissue gene expression, and tissue protein concentrations after our two-hit schedule were analyzed using a two-way ANOVA using DSS and DSP-4 as the two between-subjects factors. A three-way ANOVA was conducted for gene expression analyses in brain that included sex as a third between-subjects factor. *Post hoc* testing was done with Tukey’s correction using the Emmeans package, and CLDs were generated using the Multcomp package based on the contrasts run by Emmeans. All data visualizations were created using the ggplot2 syntax.

## Results

### DSP-4 is selective for NE and its action is reversible in the periphery

First, we sought to confirm the specificity of DSP-4 for brain NE, as its effect in the gastrointestinal system has not yet been examined. At 7 days post-injection (dpi), NE was depleted significantly by about 60% by DSP-4 in the cortex, while DA and 5HT were unaffected (Fig. 1a). However, colonic NE was also depleted by about 25% at 7dpi, while DA and 5HT were again unaffected in the colon (Fig. 1b), suggesting that the DSP-4 effect was not limited to the brain at 7dpi. Although, it has been suggested that the effects of DSP-4 are reversible in the periphery. Indeed, we found that NE concentrations in the colon returned to baseline levels after 28dpi, while cortical NE remained depleted by over 50% during the same timeframe (Fig. 1c). We also found a generalized brain loss of NET+ fibers (Fig. 1d) and a visible decrease in DBH+ cells in the LC (Fig. 1e) by 3D immunolabeling with iDISCO+ and fluorescence light-sheet microscopy. Meanwhile, colonic NE innervation, visualized by TH+ (Fig. 1f) and NET+ (Fig. 1g) fibers in the proximal colon, appeared unaffected by DSP-4. Thus, DSP-4 was selective for NE, and this effect on NE was persistent in the brain.

**Figure 1:**
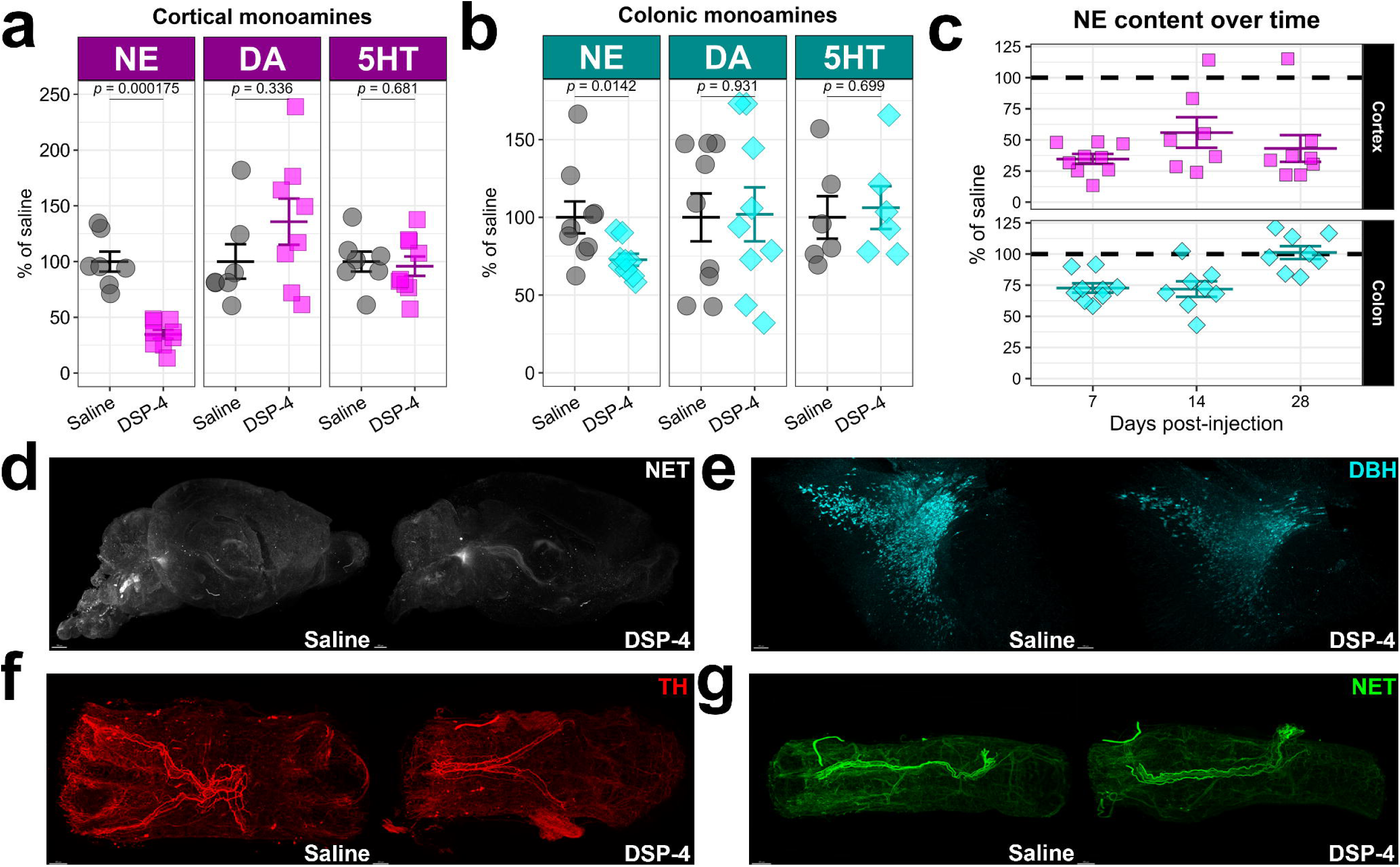
DSP-4 is selective for NE and its action is reversible in the periphery. (a) Concentrations of cortical norepinephrine (NE), dopamine (DA) and serotonin (5HT) measured by HPLC, expressed as a percentage of the average saline-treated concentration. (b) Concentrations of colonic norepinephrine (NE), dopamine (DA) and serotonin (5HT) measured by HPLC, expressed as a percentage of the average saline-treated concentration. (c) Concentrations of NE in cortex and colon at several time-points post-DSP-4 injection. The horizontal dashed line marks the concentration of NE in vehicle-treated tissues. (d) Mouse hemibrains immunostained for the norepinephrine transporter (NET) with iDISCO at 7dpi. (e) Mouse hindbrains immunostained for dopamine beta-hydroxylase (DBH) with iDISCO at 7dpi. (f) Mouse colons stained for tyrosine hydroxylase (TH) with iDISCO at 7dpi. (g) Mouse colons stained for NET with iDISCO at 7dpi. All data are presented as mean +/− SEM.

### DSP-4 does not interfere with the induction of colitis with DSS

To further ensure DSP-4’s specificity and verify that the colon’s response to injury is unaffected by peripheral DSP-4 administration, we pre-treated mice with DSP-4 or saline vehicle and subsequently subjected them to DSS-induced colitis 28dpi. The outcomes of a DSS-induced colitis schedule designed to induce peak colonic injury and inflammation (Fig. 2a) were confirmed not to be affected by DSP-4, as reflected by the lack of differences in weight loss during colitis and no obvious differences in disease severity with or without DSP-4 (Fig. 2b-c). Gross indices of disease, including increases in spleen weight (Fig. 2d) and colon shortening (Fig. 2e) after DSS were also unaffected by DSP-4. Similarly, a colitis schedule designed to induce peak neuroinflammation (Fig. 2f) was not affected by DSP-4, again as reflected by the lack of differences in weight loss (Fig. 2g), disease severity (Fig. 2h), spleen weight (Fig. 2h), or colon length (Fig. 2i) with or without DSP-4.

**Figure 2:**
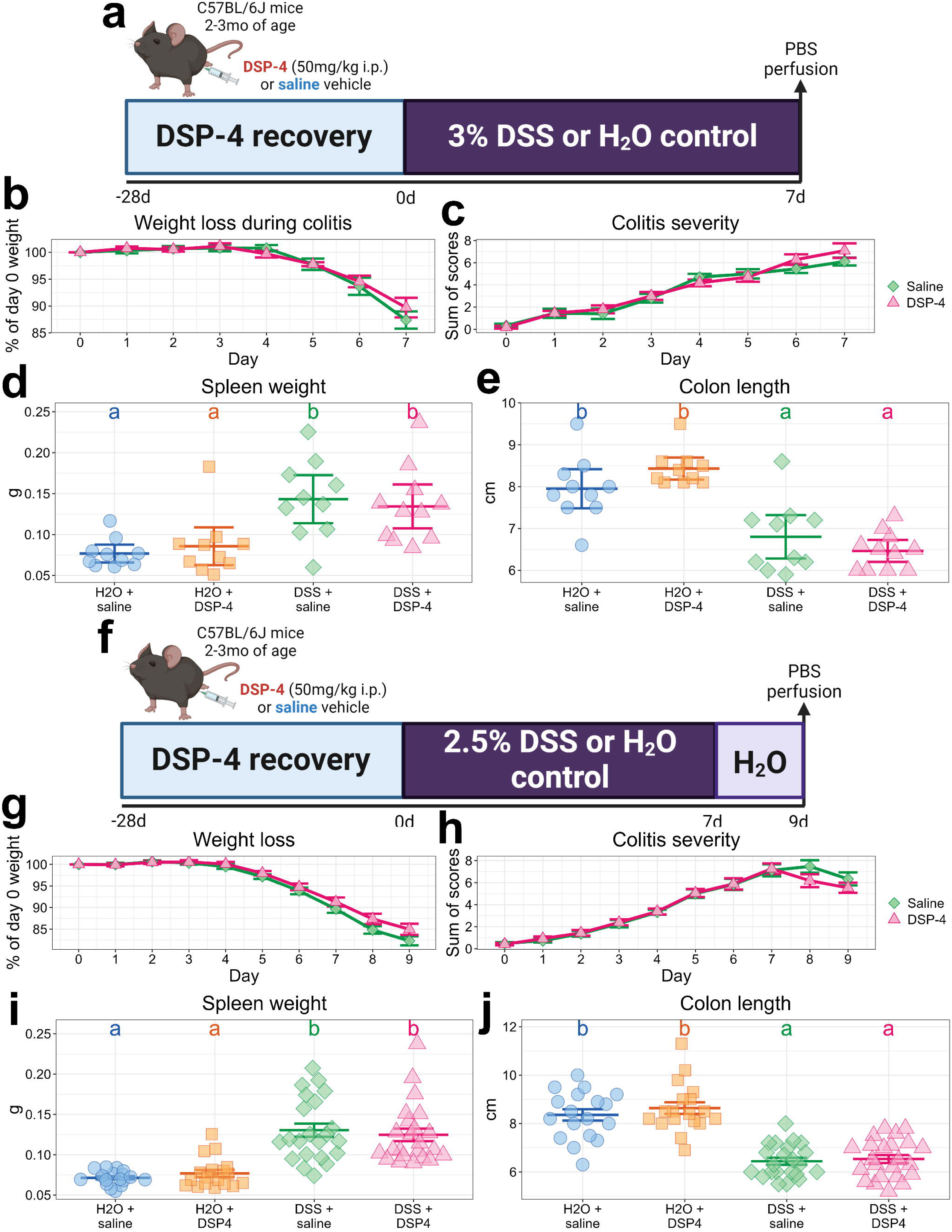
LC injury with DSP-4 does not affect the robustness of DSS-induced colitis. (a) DSS dosing schedule designed to elicit maximal gut injury. (b) Weight loss during the schedule in (a), expressed as a percentage of baseline. (c) DAI scores during the schedule from (a). (d) Spleen weight in grams after the schedule in (a). (e) Colon length in centimeters after the schedule in (a). (f) DSS dosing schedule designed to elicit peak neuroinflammation. (g) Weight loss during the schedule in (f), expressed as a percentage of baseline. (h) DAI scores during the schedule from (f). (i) Spleen weight in grams after the schedule in (f). (j) Colon length in centimeters after the schedule in (f). All data are presented as mean +/− SEM. Letters above groups denote significance from *post hoc* testing, where groups that share a letter are not statistically significantly different (*p* > 0.05).

Next, we examined the extent to which peripheral DSP-4 administered prior to colitis could modify the DSS-induced colitis phenotype. Using the DSS schedule designed to cause peak gut injury (Fig. 2a), we found that ZO-1 protein levels were not affected in any group as measured by immunoblotting (Fig. 3a-b), but the mRNA encoding for ZO-1 was downregulated by DSS treatment without being modified by DSP-4 (Fig. 3c). Similarly, the abundance of the tight junction protein occludin was not affected significantly by DSS (Fig. 3d-e) but the transcript was significantly depleted by DSS without being modified by DSP-4 (Fig. 3f). The tight junction protein claudin-5 was significantly affected by DSS but not DSP-4 (Fig. 3g-h). In total, gut barrier disintegration phenotypes induced by DSS were not modified by prior administration of peripheral DSP-4.

**Figure 3:**
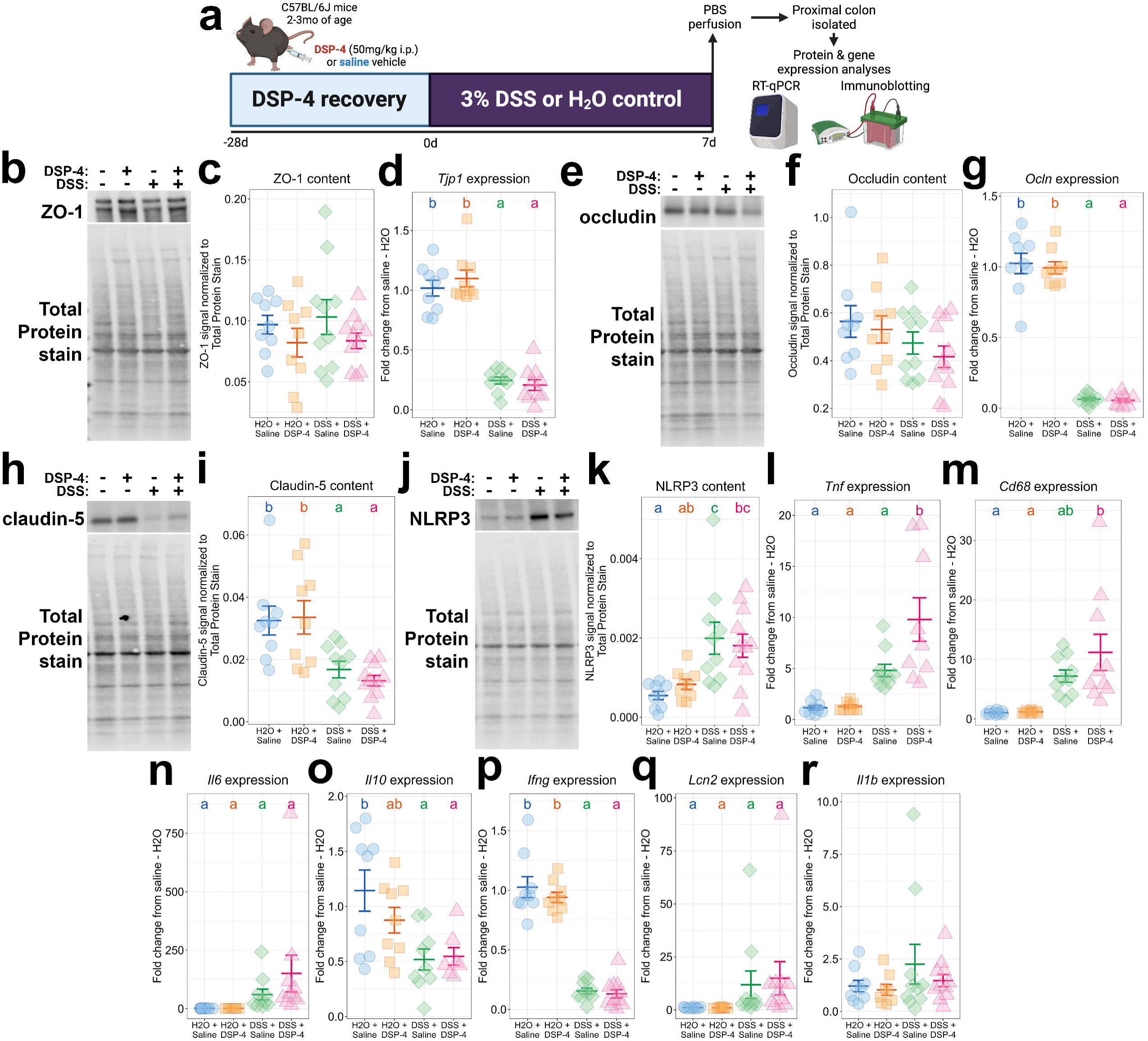
Pre-administration of DSP-4 does not affect gut barrier injury or gut inflammation in the DSS model. (a) Experimental workflow. (b) Representative western blot of ZO-1 content in the proximal colon, quantified in (c). (d) *Tjp1* expression in proximal colon measured by qPCR. (e) Representative western blot of occludin in the colon, quantified in (f). (g) Colonic *Ocln* expression measured by qPCR. (h) Representative western blot of claudin-5 in the proximal colon, quantified in (i). (j) Representative western blot of colonic NLRP3, quantified in (k). The expression of *Tnf* (l), *Cd68* (m), *Il6* (n), *Il10* (o), *Ifng* (p), *Lcn2* (q), and *Il1b* (r) in the proximal colon was measured by qPCR. All data are presented as mean +/− SEM. Letters above groups denote significance from *post hoc* testing, where groups that share a letter are not statistically significantly different (*p* > 0.05).

We also examined the extent to which DSP-4 affected colonic inflammation induced in the DSS model. The NLRP3 inflammasome, the assembly that creates mature IL-1B, was induced by DSS in the colon similarly across DSP-4 and vehicle groups (Fig. 3i-j). We observed a significant interaction between DSS and DSP-4 in the induction of *Tnf* transcript expression in the proximal colon (Fig. 3k), but the expression of *Cd68* (Fig. 3l), *Il6* (Fig 3m), *Il10* (Fig. 3n), *Ifng* (Fig. 3o), and *Lcn2* (Fig. 3p) were all modulated significantly by DSS but not DSP-4. Meanwhile, *Il1b* expression was not affected in any group (Fig. 3q). In sum, peripheral DSP-4 administration prior to colitis did not modify DSS-induced colonic inflammation in the DSS model. Overall, DSP-4’s specificity for the central compartment, namely after 28dpi when the colonic NE depletion is reversed, enables one to confidently combine DSP-4 NE denervation with other peripheral inflammatory insults. In this case, DSS-induced colitis phenotypes remain intact despite the prior peripheral DSP-4 administration, permitting unconfounded interpretations of the combinatorial effects of gut inflammation and brain NE depletion on neuroinflammation.

### DSP-4 modulates neuroimmune gene expression in the midbrain during DSS-induced colitis

To determine whether LC injury and increased gut permeability interact to affect neuroinflammation in the midbrain, we first used qPCR to measure the expression of key cytokines and chemokines after a DSS-induced colitis schedule designed to elicit peak neuroinflammation (Fig. 4a). Interestingly, the induction of *Il1b* expression in the midbrain was achieved only in mice subjected to both injuries, and this appeared to be a male-specific effect (Fig. 4b). The induction of *Ccl2* expression in the midbrain followed a similar pattern, showing an increase only in males treated with both DSS and DSP-4 (Fig. 4c). In contrast, *Tnf, Lcn2*, and *Nfe2l2* were activated by DSS without any significant effect of DSP-4, mostly in male mice (Fig. 4d-f).

**Figure 4:**
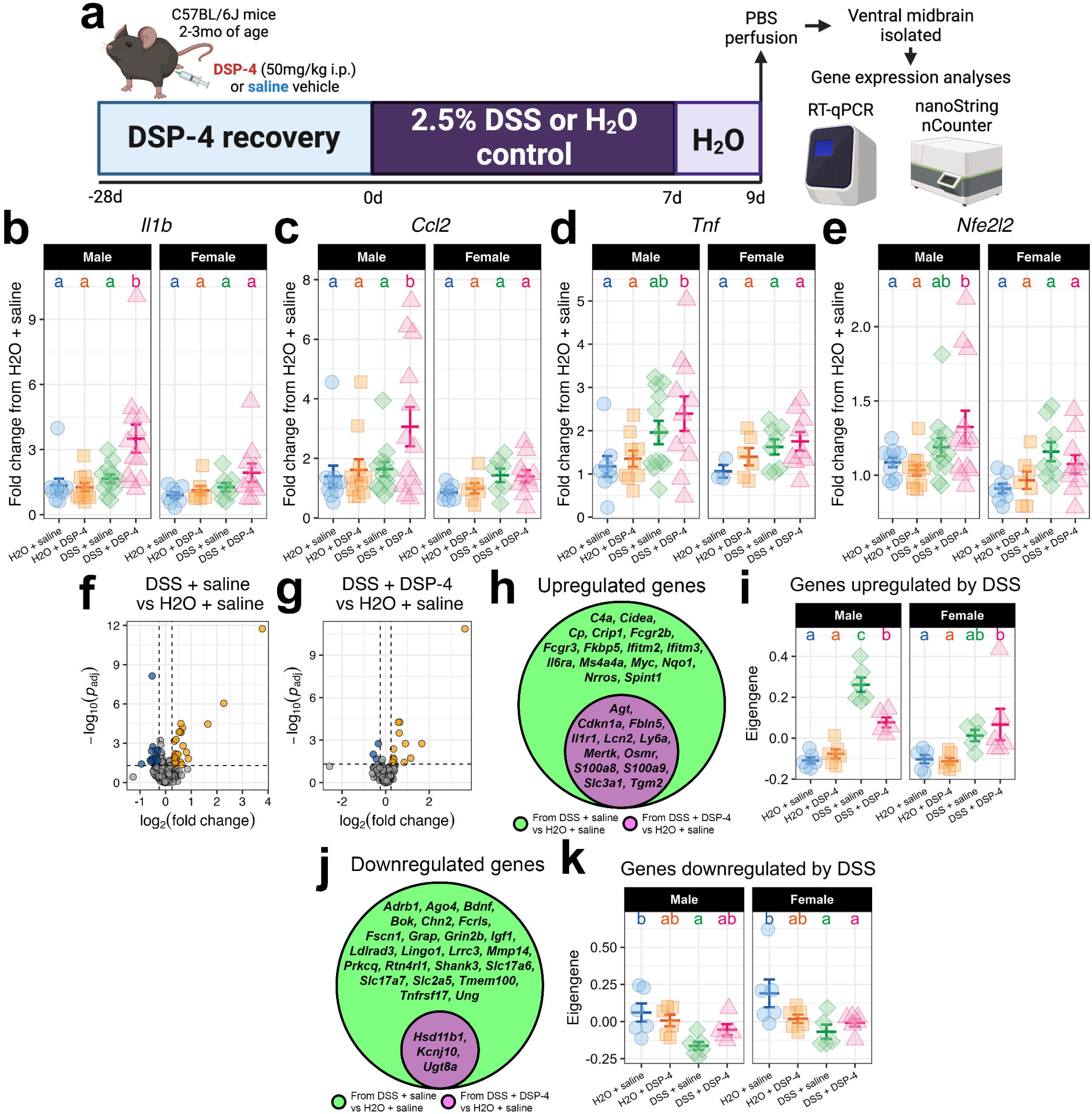
LC injury modifies midbrain neuroimmune and antioxidant gene expression responses in males triggered by gut inflammation. (a) Experimental workflow. The midbrain expression of *Il1b* (b), *Ccl2* (c), *Tnf* (d), and *Nfe2l2* (e) was measured with qPCR. (f) Volcano plot of DEGs in the DSS + saline condition relative to the double vehicle condition. (g) Volcano plot of DEGs in the DSS + DSP-4 condition relative to the double vehicle condition. For (f-g), dashed lines indicate cutoff values for labeling a gene as a DEG, and gold points reflect upregulated genes while blue points reflect downregulated genes. (h) Venn diagram listing the upregulated genes in the DSS + saline condition (green) and in the DSS + DSP-4 condition (magenta). (i) The eigengene of a gene module consisting of all genes listed in (h) was computed and plotted by condition. (j) Venn diagram listing the downregulated genes in the DSS + saline group (green) and in the DSS + DSP-4 group (magenta). (k) The eigengene of a module consisting of all genes listed in (j) was computed and plotted by condition. For (b-e), (i), and (k), data are presented as mean +/− SEM. Letters above groups denote significance from *post hoc* testing, where groups that share a letter are not statistically significantly different (*p* > 0.05).

To extend these findings, we employed the Mouse Neuroinflammation panel on the NanoString nCounter direct transcript counting platform with their Mouse Neuroinflammation panel with an additional 55 custom genes. Standard differential expression analyses revealed 53 differentially expressed genes (DEGs) in the midbrains of DSS-only mice and 15 DEGs in the midbrains of mice that received both insults, relative to vehicle-treated mice (Fig. 4f-g). Of the upregulated genes in both conditions, none were unique to the two-hit group; DSS-only mice upregulated all the genes upregulated by the two-hit group, including *Cdkn1a*, *Il1r1*, *Ly6a*, *Lnc2*, and *S100a9*, as well as unique genes such as *C4a*, *Fcgr3*, *Il6ra*, and *Nrros* (Fig. 4h). Examining the first principal component of the genes upregulated by DSS alone revealed that DSS-treated mice expectedly upregulate this module, while mice pre-treated with DSP-4 show a blunted response in males only (Fig. 4i). DSS had a smaller but similarly positive effect on these genes in female midbrains, and DSP-4 did not influence this upregulation (Fig. 4i). Similarly, of the downregulated genes in both conditions, none were unique to the two-hit group; mice treated with both DSS and DSP-4 downregulated only *Hsd11b1*, *Kcnj10*, and *Ugt8a*, while DSS-only mice downregulated these genes as well as *Adrb1*, *Bdnf*, *Fcrls*, *Lrrc3*, *Shank3*, *Tnfrsf17*, and others (Fig. 4j). The eigengene of this module was expectedly downregulated by DSS alone but not by DSS and DSP-4 together in males, while female mice downregulated this module in both conditions similarly but to a lesser extent than males (Fig. 4k). Taken together, these data suggest that LC injury modulates the midbrain neuroimmune response to intestinal permeability. LC injury activates the expression of specific cytokines and chemokines in the midbrain, including *Il1b* and *Ccl2*, which are normally absent during colitis, and blunts the differential expression of genes normally associated with colitis.

## Discussion

In this study, we established and characterized a novel two-hit mouse model that incorporates several common and early features of PD, including gastrointestinal inflammation and LC degeneration. Specifically, we provide a reevaluation of the DSP-4 neurotoxin model of LC injury, confirming its specificity for NE as expected based on previous studies^76–79^. We also show gastrointestinal NE depletion via DSP-4 is reversible after 4 weeks, affirming early reports of transient sympathetic denervation with DSP-4^77,79,80^. Importantly for our study, the peripheral reversibility of DSP-4 meant that the induction of colitis via DSS and the colonic features of DSS-induced colitis were left intact. This feature of our two-hit model is critical in the downstream interpretations of any effects observed on ventral midbrain outcomes. Several previous studies have employed pharmacological or genetic tools, such as anti-TNF therapies, paquinimod, flavonoids, and targeted *LRRK2* mutations^15,19,20,81^, to modulate the brain effect of the colitis. In the DSS model, our group has previously demonstrated that colitis severity is the best predictor of neuroinflammation^14^, and modifying the severity of the disease modifies the neuroinflammatory phenotype^82^. Those efforts, while still critical for the field, affect the induction of colitis itself, making it difficult to determine whether neuroinflammation itself is modified or whether the systemic insult that triggers this neuroinflammation is modified. Here, we show that a brain-restricted phenotype, LC injury, affects the neuroinflammatory response in the midbrain induced by a leaky gut.

We observed that *Il1b* expression is significantly activated in the ventral midbrain of mice that received both LC injury and colitis but not in mice that received either insult alone. Previous studies support an interaction between NE and IL-1β during inflammatory challenges. *In vitro* studies have demonstrated that lipopolysaccharide (LPS) induces IL-1β production in microglia, which can be blocked with the co-application of NE^56^. NE may activate suppressors of IL-1β signaling, including IL-1ra and IL-1RII^83^, or suppress the expression of *Il1b* via the suppression of the nuclear action of NF-κB^84^, which may downregulate interferon regulatory factor 1 (IRF1)^85^, a transcription factor that activates *Il1b*. IL-1β overproduction is thought to be a driver of PD, as IL-1β induction has been shown to worsen motor dysfunction and alpha-synuclein pathology in animal models of PD-like pathology^86^, and circulating IL-1β has been associated with worsened PD severity^87,88^. As such, the observed activation of *Il1b* expression in the ventral midbrain of mice treated with DSS only in the presence of an injured LC suggests potentially damaging consequences for the nigrostriatal pathway.

Similarly, we found selective activation of *Ccl2* expression in the ventral midbrain of mice that received both DSS and DSP-4. CCL2 is a major chemoattractant implicated in the infiltration of myeloid cells, especially monocytes and neutrophils, into the brain during inflammation^89,90^. A relationship between NE and CCL2 has not been extensively examined in microglia, but a recent report demonstrated that NE reduces CCL2 production in mouse and human macrophages in a β-adrenergic receptor-dependent manner^91^. Additionally, recent work in astrocytes has revealed a complicated relationship between NE and CCL2 that may provide insight for the data in this study and others. At rest, NE induces CCL2 production in astrocytes, likely through β-adrenergic receptors^92,93^, while clonidine, an α2-adrenergic receptor agonist, reduces CCL2 production in these cells^94^. Further, endotoxin-exposed astrocytes downregulated CCL2 with NE^95^, suggesting that the immunoregulatory effect of NE depends on the neuroinflammatory milieu. In all, our data are in line with this notion; the expression of *Ccl2* was induced by NE depletion via DSP-4 only during peak colitis. This effect raises the interesting prospect of peripheral immune cell invasion into the brain, which future studies will address. The influx of CCR2+ peripheral phagocytes has been reported in PD and mouse models of PD^24,96^, suggesting that the induction of their ligand CCL2 is a relevant event in our model, and dysregulation of the crosstalk between brain and peripheral immune system is a potential consequence of NE depletion.

We found that the differential regulation of several genes during DSS-induced colitis was dampened by LC injury. At first glance, this may seem to suggest that LC degeneration is neuroprotective during colitis, but closer inspection of their identity and their associated pathways reveals that upregulation of several ventral midbrain genes (*Ms4a4a, Fcgr2, Myc, Lcn2, Osmr, Nqo1, and Nrros*) is likely to represent a protective neuroinflammatory and antioxidant response. For example, the rs1582763 variant in the *MS4A* locus increases *MS4A4A* expression^97^, whose mouse ortholog *Ms4a4a* was upregulated by DSS but not DSS and DSP-4 together. This *MS4A* variant was recently shown to be associated with reduced risk of Alzheimer’s disease^98^ by promoting healthy lipid metabolism and reducing chemokine signaling in microglia^97^. FcγRIIB is the only inhibitory Fc receptor, known for controlling many immune processes and regulating a healthy defense against infection^99^, and the murine gene encoding for this Fc receptor *Fcgr2* was upregulated by DSS-induced colitis alone but not when the LC was injured in mice treated with both DSS and DSP-4. Similarly, MYC, encoded by *Myc* which was upregulated by DSS alone in our study, has been shown to control inflammatory activity in myeloid cells^100^. Lipocalin-2, encoded by *Lcn2*, and oncostatin M signaling through its receptor, encoded by *Osmr*, which are both upregulated during DSS irrespective of DSP-4 treatment, are thought to be highly neuroprotective during inflammation^101–104^. Finally, DSS-only mice upregulated *Nqo1* and *Nrros*, genes encoding for critical regulators of antioxidant responses^105–107^, while mice given both DSS and DSP-4 did not. Together, we conclude that many of the DSS-associated genes identified here are protective against inflammatory and oxidative stress, and the DSP-4-associated ablation of their activation would be expected to be damaging to the nigrostriatal pathway.

At the same time, some genes regulated by DSS treatment may be injurious components of the neuroimmune response to colitis. Genes like *Il1r1*, *Cdkn1a*, and *S100a9* were upregulated by DSS in both the presence and absence of DSP-4. S100A9 activates NF-κB signaling and has been shown to aggravate neuroinflammation in a model of subarachnoid hemorrhage^108^ and Alzheimer’s disease^109^ and has been implicated as an integral component of inflammation along the gut-brain axis in the DSS model^15^. Cyclin-dependent kinase inhibitor p21, encoded by *Cdkn1a*, appears to be an inflammatory response element and may be a marker of DNA damage^110,111^, and an upregulation of the IL-1 receptor type 1, encoded by *Il1r1*, has been associated with pathogenesis in several neurodegenerative diseases^112^. In general, the colitis-induced neuroimmune response is complex, and further work will be needed to unravel this complexity especially amidst LC injury.

We acknowledge that a notable limitation of our study is our inability to determine whether the detectable regulation of neuroimmune gene expression in the ventral midbrain actually compromise dopaminergic neuron survival. To address this, future studies will employ a chronic dosing scheme of DSS to induce colitis after LC injury, that has been used previously to induce neurodegeneration^19,20,113,114^. Another step in this line of investigation might be to investigate *in vitro* the regulation of genes identified by our *in vivo* study, using NE or adrenergic receptor drugs amidst LPS treatment of microglia or astrocytes, as done previously^56,94^, and/or by mining the ever-growing body of high-throughput sequencing studies of neurodegenerative diseases and their respective mouse models. However, based on the existing literature demonstrating that an LC lesion worsens other models of neuroinflammation and degeneration^58,60,61,63–66^, we predict that the dampened ventral midbrain neuroinflammatory response to DSS-induced colitis we report here due to LC injury will have a detrimental outcome for the nigrostriatal pathway, which has been shown by numerous studies to be selectively vulnerable to the effects of inflammatory and oxidative stress^24,115–117^.

Another limitation of our study is that we cannot discern whether the changes in neuroimmune gene expression observed here are due to the depletion of NE or another transmitter from the LC. To elucidate the role of NE, future studies would combine our two-hit model with pharmacological drugs for adrenergic receptors or NE reuptake and examine the effects on neuroinflammation. This strategy has been employed in several mouse studies previously; two NE reuptake inhibitors, such as desipramine and atomoxetine, and β2-adrenergic receptor agonists, such as clenbuterol, have lessened neuroinflammation after immune insults and alpha-synuclein overexpression ^118–120^, while β-adrenergic receptor blockers, like metoprolol and propranolol, have exacerbated aging– and proteinopathy-related neuroinflammation and subsequent cognitive dysfunction^54,121^. Based on these data, we expect that the loss of NE mediates most of the inflammatory phenotype we observe in our two-hit model, however the anti-inflammatory effects of brain-derived neurotrophic factor^122^ and galanin^123^, some of the LC’s major co-transmitters, are also worth further consideration.

With regards to the relevance of our findings within the context of therapeutic interventions, identifying the molecules mediating the inflammatory predisposition that comes with LC injury may inform therapeutic strategies for PD. NE medication is routinely used in the treatment of PD. For example, propranolol is used to treat tremor^124^ and control levodopa-induced dyskinesia in PD^125^, desipramine and other norepinephrine reuptake inhibitors are used to treat depression and anxiety in PD^126–129^, and the NE reuptake inhibitor atomoxetine is being examined as a therapy for cognitive decline in PD^130,131^. Given the role of NE signaling in brain health and immunity, the effect of these drugs on neuroinflammation must be considered. In this vein, a recent phase II trial examining the efficacy of atomoxetine in protecting cognition in Alzheimer’s disease demonstrated that NE reuptake inhibitor treatment reduced phosphorylated tau and CD244, a putative marker for cytotoxic T-cells and natural killer cells, in the cerebrospinal fluid ^132^. Including readouts of systemic and central inflammation in further work studying these therapeutics in PD may build further evidence that NE signaling is protective against neuroinflammation-associated neurodegeneration.

In summary, we have developed a two-hit mouse model that models common and disease-modifying features of early PD. With this model, we have observed that DSS-induced colitis triggers what is likely a compensatory neuroimmune gene program in the ventral midbrain, while activating very little expression of pro-inflammatory cytokines and chemokines. When the LC is injured, these programs are reversed and potentially protective immune and antioxidant responses are dampened. Overall, this study highlights the LC as a regulator of neuroinflammation during colonic permeability and inflammation. This study, alongside the mounting evidence positing the LC as neuroprotective, may inform future therapeutic strategies involving NE drugs to prevent or delay PD progression and draw focus to LC integrity as a predictor of disease progression.

## Figures

**Figure S1:**
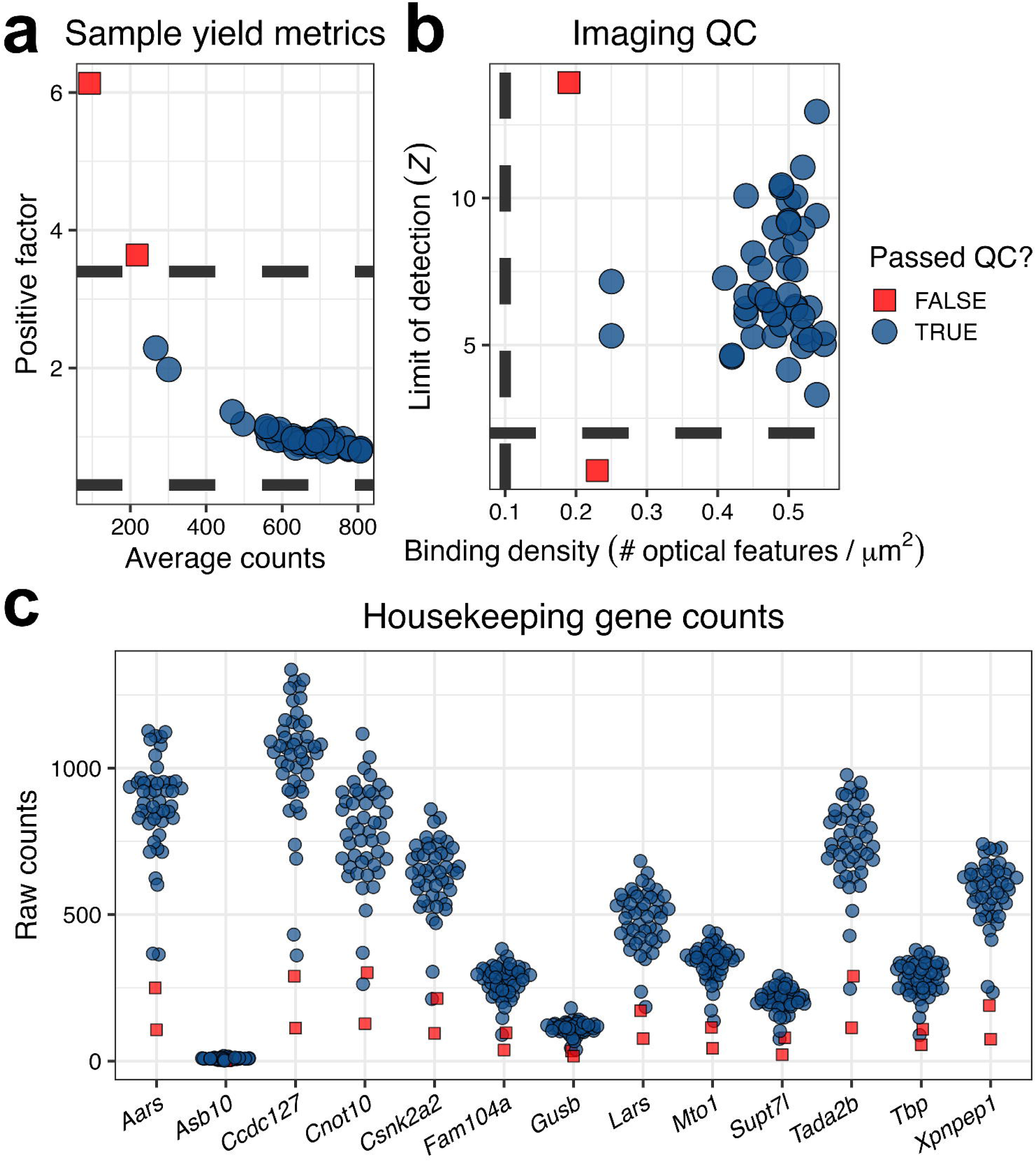
Quality assessment of NanoString nCounter data. (a) Sample yield metrics, with the average transcript count for each sample on the x-axis and the positive library scaling factor based on the positive control probes on the y-axis. Dashed lines mark the lower and upper cutoffs for the positive factor. (b) Imaging quality metrics, with the binding density (expressed as the number of features per square micron) on the x-axis and the limit of detection on the y-axis. Dashed lines mark the lower bounds for each metric. (c) Raw counts of all housekeeping genes included in the panel. From this figure, *Asb10* was omitted as a housekeeper due to its low counts overall. For each panel, each dot represents a sample, and samples are colored by whether they pass all QC checks and were thus carried into downstream analyses.

## Author’s contributions

JSB, OUH, and MGT contributed to the overall design and organization of the research. JSB, OUH, JH, CLC, and NKN contributed to the execution of animal studies, including day-to-day animal care and euthanasia. JSB, CLC, and NU performed biochemical experiments. JSB analyzed data and created figures. JSB, OUH, and MGT wrote and edited the manuscript. All authors reviewed and approved the final draft of the manuscript.

## Conflict of interest

The authors declare no conflicts.

## Acknowledgements

We thank members of the Tansey lab for useful discussions, insight, and feedback concerning this study. Figures 2-4 were created using BioRender.com. Partial funding for this work was derived from awards from the NIH T32-NS082128 (JSB), NIH NIA 1RF1AG057247 (MGT), NIH NINDS 1RF1NS28800 (MGT), and the joint efforts of The Michael J. Fox Foundation for Parkinson’s Research (MJFF) and the Aligning Science Across Parkinson’s (ASAP) initiative. MJFF administers the grant ASAP-020527 on behalf of ASAP and itself. For the purpose of open access, the author has applied a CC-BY public copyright license to the Author Accepted Manuscript (AAM) version arising from this submission.

## Notes

### Competing Interest Statement

The authors have declared no competing interest.

